# An Internal Model of Sensorimotor Context in Freely Swimming Electric Fish

**DOI:** 10.1101/2022.06.08.495343

**Authors:** Avner Wallach, Nathaniel B. Sawtell

## Abstract

Nervous systems are hypothesized to learn and store internal models that predict the sensory consequences of motor actions. However, little is known about the neural mechanisms for generating accurate predictions under real-world conditions in which the sensory consequences of action depend on environmental context. Using novel methods for underwater neural recording in freely swimming electric fish, we demonstrate that complex movement-related input to the active electrosensory system is effectively cancelled, despite being highly-dependent on the nearby environment. Computational modeling and closed-loop electrophysiological experiments indicate that the cerebellum-like circuitry of the electrosensory lobe generates context-specific predictions of self-generated input by combining motor signals with electrosensory feedback. These results provide mechanistic insight into sophisticated internal models supporting natural behavior in freely moving animals.

**One-Sentence Summary:** Underwater recordings in electric fish reveal neural mechanisms for predicting the sensory consequences of behavior under natural conditions in freely moving animals.

## Main Text

Converging lines of evidence from theoretical, human behavioral, clinical, and neuroimaging studies suggest that the nervous system contains internal models that predict sensory input or changes in state resulting from motor commands (*1-5*). Although the neural mechanisms for internal models remain largely unknown, progress has been made by examining how sensory systems distinguish between self-generated (reafferent) and externally-generated (exafferent) sensory input (*6-9*). Copies of motor commands, known as efference copy or corollary discharge, are hypothesized predict and cancel self-generated sensory input (*10, 11*). However, for natural behavior in freely moving animals, the relationship between motor commands and reafferent sensory input is rarely fixed, but rather depends on environmental context. Patterns of optic flow on the retina, for example, depend not only on self-motion but also on the external visual scene (*12*). Such observations raise questions regarding the sufficiency of purely motor-based strategies for reafference cancellation in natural environments.

Here we examine internal models for reafference cancellation in electric fish. Prior work has elucidated the cellular and circuit mechanisms for predicting and cancelling self-generated electrosensory input at the first central processing stage in the electrosensory lobe (ELL) (*13*). Principal cells of the ELL integrate electrosensory input from electroreceptors on the skin with motor corollary discharge signals conveyed by a granule cell-parallel fiber system similar to that of the cerebellum (*14*). Closed-loop electrophysiological studies have shown that anti-Hebbian synaptic plasticity shapes granule cell synaptic input into highly-specific predictions (termed negative images) of self-generated sensory input evoked by the fish’s electric organ discharge (EOD) pulse (*15-18*). Importantly, however, past studies of reafference cancellation in electric fish have been conducted in immobilized preparations using artificial stimuli. Under natural conditions, the active electrosensory system faces the daunting challenge of detecting and characterizing small perturbations of the fish’s electrical field due to nearby objects amidst much larger perturbations due to the fish’s own movements (*19, 20*). Tail motion, for example, alters the location of the electric organ relative to electroreceptors on the skin. Moreover, electrosensory reafference due to tail motion may depend on environmental features such as the location of the fish relative to non-conducting boundaries (e.g. the river bed or the water surface).

To examine whether the ELL is capable of predicting and cancelling self-generated input under naturalistic conditions (**Fig. 1A**), we implanted custom-designed, waterproofed moveable microwire electrode arrays into the medial zone of the ELL (see **Fig. S1**). We then performed uninterrupted behavioral and neural recordings (30-134 hrs) as the fish swam freely in a circular tank (**Fig. 1B**). Overhead videography and automated pose-estimation, along with a head-mounted accelerometer, were used to track the fish’s behavior (**Fig. 1C**, see **Movie S1**). Since the ELL is somatotopically organized, each electrode recorded neural activity associated with activation of electroreceptors located on a specific region on the fish’s skin (the receptive field, RF, **Fig. 1D**; **Fig. S1**). On most electrodes, we recorded both prominent (> 1 mV) short-latency (2-6 ms) EOD-evoked local field potentials (LFPs) and single-unit spiking activity. The LFP amplitude reflects synaptic activation of the ELL by electroreceptor afferents and grades smoothly with the strength of the electrosensory input to the local RF on the skin (*21, 22*). Importantly, this approach allows for a quantitative examination of the input-output transformation performed by the ELL under freely behaving conditions in which patterns of sensory input are controlled by the behavior of the fish (rather than the experimenter).

**Fig. 1.**
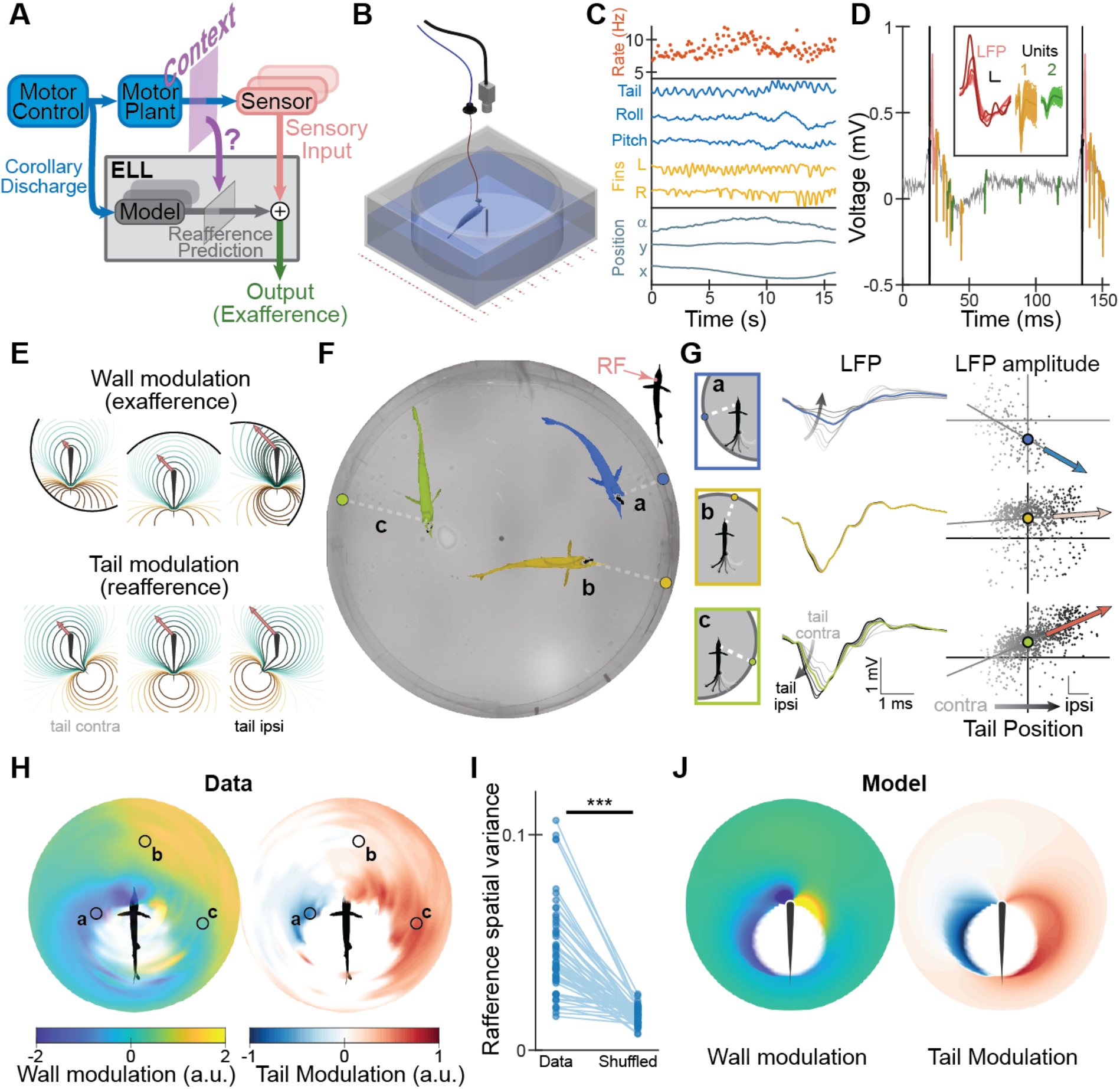
Electrosensory reafference depends on environmental context. (**A**) ELL uses motor signals to predict and cancel reafference, although it is unknown how predictions are adapted to changing sensorimotor context during natural behavior. (**B**) Arena design. (**C**) Tracked behavioral variables, from top: EOD rate; tail position, pitch/roll angles, fin positions, location in tank (α-azimuth, y, x). (**D**) Extracellular voltage trace recorded in a freely-swimming fish. Black: EOD artifact; pink: LFP; green/yellow: waveforms of two isolated single-units. Inset scale bars: 1 ms, 0.1 mV. (**E**) Model showing wall induced (exafference, top) and tail induced (reafference, bottom) sensory modulation; colored contours: electric potential; arrows: electrosensory input (transdermal potential) at left side of face. (**F**) Example fish positions; colored circles indicate wall location closest to fish. (**G**) Left: LFP traces for different tail positions; grey: contralateral to the receptive field, black: ipsilateral; colored: straight. Right: linear regression of LFP amplitude vs. tail position; colored circles: intercept; colored arrows: slope; scale-bars: z-score. (**H**) LFP regression intercept (wall modulation, left) and slope (tail modulation, right) maps. (**I**) Spatial variance of tail modulation map compared with position-shuffled control (p=7.54*10^−11^, Wilcoxon signed-rank, n=37). (**J**) Electrostatic field model replicates wall (left) and tail (right) modulation maps.

We first characterized the electrosensory input into the ELL as the fish swam in the tank. The electrically non-conductive tank wall distorts the electric field generated by the EOD, creating an externally-generated sensory modulation (exafference) that depends on the wall’s position relative to the fish (**Fig. 1E**, top). Roles for the active electrosense in navigation based on large environmental features, such as tank walls, have been demonstrated previously (*23*). Tail motion during swimming displaces the electric organ and therefore distorts the field as well, producing a self-generated sensory modulation (reafference, **Fig. 1E**, bottom). To quantify the sensory effects arising from these two sources, and their possible interactions, we used linear regression analysis of the measured LFP amplitude against the tail position *for each relative wall position* (**Fig. 1F,G**, see **Methods, Movie S2**). Performing this analysis on long stretches of continuous recording (∼7 hours or 300,000 EODs for **Fig. 1H**) allowed us to produced two maps, depicting the regression parameters as functions of relative wall position. The regression *intercept* parameter quantifies the average sensory input for a neutral (straight) tail, and thus its map reflects the sensory modulation due to the wall (i.e., the exafference component, **Fig. 1H, left**). The regression *slope* parameter describes the relationship between the sensory input and the tail’s position, and thus its map depicts the sign and magnitude of the self-generated sensory modulation (i.e., the reafference component, **Fig. 1H, right**).

The wall modulation exhibited a dipolar pattern (**Fig. 1H, left**): sensory input was attenuated when the wall faced the receptive field (e.g., position ‘a’ in **Fig. 1**) and was amplified when the wall faced the opposite side of the fish. Importantly, the tail modulation map (**Fig. 1H, right**) was spatially patterned: leftward motion of the tail (towards the receptive field) amplified the LFP in some positions (e.g. position ‘c’) while attenuating it in others (e.g., position ‘a’). Therefore, the same motor action resulted in different (and even opposite) sensory effects, depending on the environmental context (i.e. the position of the fish relative to the tank walls). A high degree of spatial patterning of the tail modulation map was consistently observed, regardless of the receptive field location (**Fig. 1I, Fig. S2**). A model of the fish’s electric field (see **Methods**) recreated both wall-induced and tail-induced LFP modulation patterns (**Fig. 1J**), demonstrating that these phenomena can be attributed to the physics of electrical fields and their interactions with the environment, rather than to internal neural processing.

The observation that the sensory consequences of motor behavior depend strongly on the environmental context raises the key question of whether and how such complex patterns of reafference are predicted and cancelled by the ELL under natural conditions. To generate hypotheses regarding how this may be accomplished, we constructed a computational model of the ELL (**Fig. 2A**), consistent with its known circuitry and synaptic plasticity (see **Methods)** and used the electric-field model (see **Fig. 1J**) to generate sensory data for all wall and tail positions from an array of receptive fields on the model fish’s body. We tested what input information is required for the model to accurately predict and cancel tail-induced reafference irrespective of the position of the fish in the tank. Cancellation was poor in models receiving only motor signals related to tail position (**Fig. 2B,C**, ‘Motor’) since they generate context-independent predictions (**Fig. 2D**). However, when the motor information was complemented by explicit context information (i.e. the position of the wall relative to the fish), nearly perfect cancelation was achieved (**Fig. 2**, ‘Motor + Context’; note in **Fig. 2D** that the internally generated tail modulation map resembles a ‘negative image’ of the input tail modulation map). Thus, cancelation of self-generated sensory input in freely behaving animals necessitates context-specific internal predictions of motor actions.

**Fig. 2.**
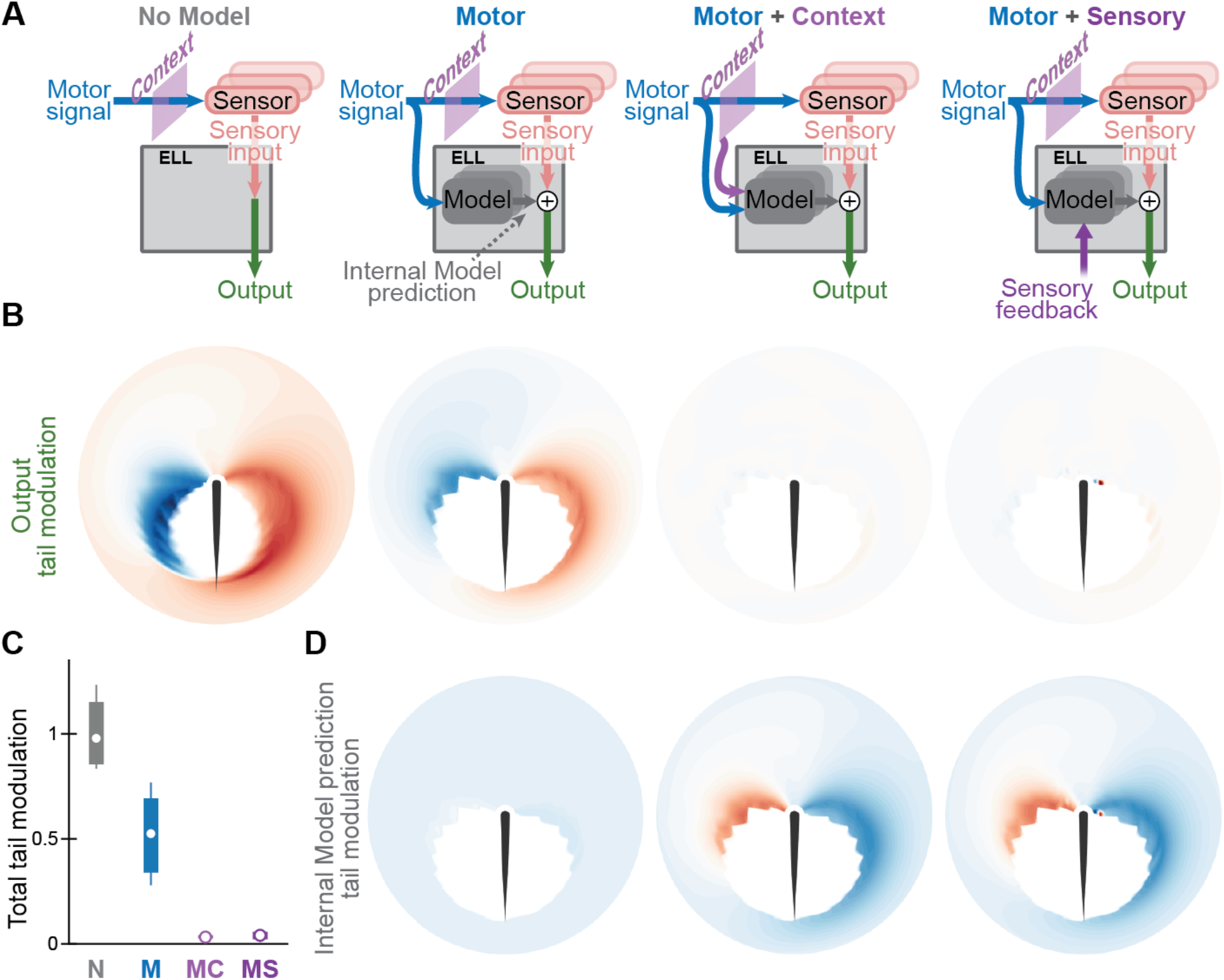
Reafference prediction and cancellation across contexts requires both motor and sensory information. (**A**) Simulated ELL internal models receiving different types of predictive information, from left: none, motor (tail position), motor and context (relative wall position), motor and sensory feedback (transdermal potential at all RFs). (**B**) Tail modulation maps (reafference) for the output (green arrow in **A**) of each ELL model. (**C**) Total tail modulation of model outputs across all RFs (normalized to average tail modulation of sensory input). N, none; M, motor; MC, motor+context; MS, motor+sensory feedback. Boxes, IQR; whiskers, 10-90 percentile range. (**D**) Tail modulation map of internal model prediction (gray arrow in **A**). Motor internal model produces spatially-uniform (context-independent) predictions; motor+context and motor+sensory produce a ‘negative image’ of the input reafference (compare to **B**, left).

How might the ELL derive information about environmental context? Large environmental features (such as the tank walls in our experiments) activate electroreceptors distributed across the body surface of the fish. In addition to motor corollary discharge and tail proprioception (*24-26*), granule cells of the ELL receive prominent electrosensory feedback from higher processing stages (*27*). Hence, individual ELL neurons could integrate electroreceptor afferent input originating from a local receptive field on the skin with granule cell-parallel fiber input conveying both motor signals related to tail movements and context information originating from spatially distributed electroreceptors located elsewhere on the body. Consistent with this hypothesis, models provided with both motor information and sensory signals from all RFs also generated context-specific reafference predictions and achieved near-perfect cancelation (**Fig. 2**, ‘Motor + Sensory’).

To test whether such context-specific cancellation is exhibited by ELL neurons in freely swimming fish, we analyzed the responses of well-isolated single-units, recorded simultaneously with LFPs. These experiments used a richer scenario in which a conductive object – a vertical brass pole – was placed at the tank’s center (**Fig. 3A,B, Movie S3**). Two distinct unit types were encountered (**Fig. 3C, Fig. S3**): type 1 units exhibited low baseline firing rates (0-6 Hz) and short-latency spiking time-locked to each EOD, while type 2 units exhibited higher baseline firing (6-25 Hz) and weaker responses to each EOD. Based on prior *in vivo* intracellular recordings from morphologically identified cells (*28, 29*), type 2 units correspond to the output cells of the ELL. Responses of type 1 units were consistent with several different interneuron and afferent fiber classes characterized previously. Importantly, our prediction from prior work is that the cancellation of self-generated sensory input will be evident at the level of the output cells of the ELL (*18, 25, 28*), i.e. in type 2 units.

**Fig. 3.**
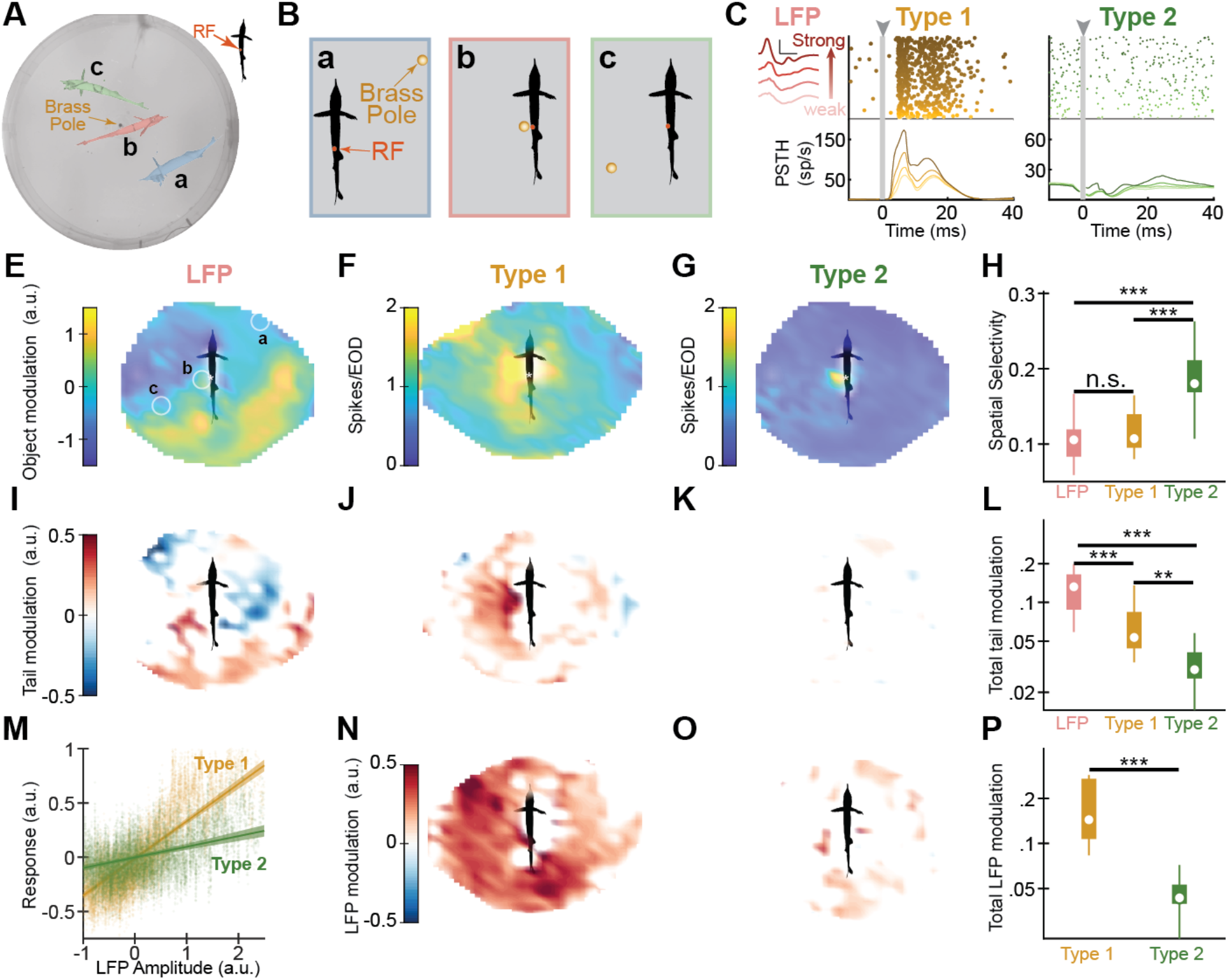
Context and movement invariant representation of external objects in the ELL of freely swimming fish. (**A**) Example fish positions with a brass pole at tank center. (**B**) Corresponding relative object positions. (**C**) EOD-aligned raster plots (top) and peri-stimulus time histograms (PSTH, bottom) for example Type 1 (putative interneurons, yellow) and Type 2 (putative output cells, green) units, ordered by concurrently measured LFP amplitude (pink, left). (**E-G**) Object modulation (exafference) map for LFP, Type 1, and Type 2 units with similar RF centers (white asterisk). Circles in (E) correspond to object positions in (B). (**H**) Object modulation spatial selectivity is significantly increased in Type 2 units (p=0.14 LFP-Type1, p<10^−5^ LFP-Type2, p=3*10^−5^ Type1-Type2). (**I-K**) Tail modulation (reafference) maps for same example units. (**L**) Significant reduction in total tail modulation from LFP to Type 2 units (log-scale. p=0.00047 LFP-Type1, p<10^−5^ LFP-Type2, p=0.004 Type1-Type2). (**M**) Overall LFP (sensory input) sensitivity of Type 1 and Type 2 units. Circles, neural response (z-scored, boxcar smoothed); bold line, linear fit; shading, 95% confidence interval. (**N-O**) Position-controlled LFP modulation map (local slope with respect to LFP amplitude) for same unit examples. (**P**) Significant reduction in total LFP modulation in Type 2 units (log-scale. p=2*10^−5^). Boxes, IQR; whiskers, 10-90 percentile range. All p-values: random permutation test, LFP n=40, Type 1 n=9, Type 2 n=26. n.s. p≥0.05, ** p<0.01, *** p<0.001.

We repeated the linear regression analysis described above but this time in coordinates of object (rather than wall) position relative to the fish. The resulting object and tail modulation maps revealed striking differences in the sensory information contained in LFPs, type 1, and type 2 units. First, the object modulation (exafference) maps of both LFPs and type 1 units were spatially diffuse, with graded activation due to both the surrounding wall and the center object (**Fig. 3E,F)**. In contrast, type 2 units (putative output cells) were highly-selective, responding only when an object was located within a spatially restricted RF (**Fig. 3G**; see additional unit examples in **Fig. S4**). This significant increase in spatial selectivity (**Fig. 3H)** cannot be explained by a simple non-linear transformation of the LFP input, such as thresholding (see **Fig. S5**). Second, while the pronounced, spatially-patterned tail modulation (reafference) observed for LFPs (**Fig. 3I**) was only slightly reduced in type 1 units (**Fig. 3J**), it was almost completely cancelled across the entire spatial map in type 2 units (**Fig. 3K,L**). Hence, putative output cells of the ELL exhibit dramatic cancellation of tail-induced reafference during free behavior, irrespective of external context.

Although our analysis thus far has focused on tail motion, additional motor variables likely contribute to the total reafferent input to the ELL. To evaluate the cancellation of total reafference in the ELL, we compared LFP and single-unit responses recorded on the same electrode. Overall, type 1 unit responses exhibited a stronger correlation with LFP amplitude than those of type 2 units (putative output cells) (**Fig. 3M**). Since LFP amplitude reflects the total sensory input to a local region of the body surface, modulations in LFP amplitude occurring when the exafference is constant (i.e. for a fixed position of the fish in the tank) provide a measure of the total reafferent input due to the fish’s behavior. By performing the linear regression analysis, described above, using the LFP amplitude as the input, we produced an *LFP modulation map* that quantifies the sensitivity of single-unit spiking to total reafference. While type 1 unit responses scaled with LFP amplitude at almost all positions (**Fig. 3N**), type 2 unit responses were far less sensitive to changes in LFP amplitude (**Fig. 3O-P**; see **Fig. S5**). These results demonstrate a dramatic transformation of sensory representations within the ELL, from a sensory input heavily dominated by reafference to a neural output conveying a behavior-and context-invariant representation of external objects.

Our modeling suggests that invariant responses to exafferent input in type 2 units arise from context-specific predictions generated by a learning process within the ELL (**Fig. 2D**). Specifically, we hypothesize that context-specific negative images of reafferent input are shaped by anti-Hebbian plasticity acting on granule cell inputs conveying both motor signals and spatially distributed electrosensory input. Testing this hypothesis in freely swimming fish is challenging, however, as it requires manipulating the correlations between sensory and motor signals and isolating their effects on neural responses. To achieve the required experimental control, we turned to an immobilized closed-loop preparation. Neuromuscular paralysis was used to block the EOD while leaving its motor command (and the associated motor corollary discharge pathway) intact. Sensory input was generated using brief electrical pulses delivered to electroreceptors on the face (inside the receptive field of recorded ELL neurons) (**Fig. 4A,B**, *Sensory Stim*). This input was made predictable (simulating reafference) by locking the stimulus to the fish’s emitted EOD motor command (**Fig. 4A,B**, *Motor command*). Context information was provided by a second electrosensory stimulus positioned over electroreceptors far from the face (i.e. outside of the receptive field of the recorded ELL neurons) (**Fig. 4A,B**, *Context Cue*). To induce plasticity and negative image formation, stimuli were delivered repeatedly in two alternating configurations: in ‘*context a’*, sensory stimuli had positive polarity and were shortly preceded by a context cue, while in ‘*context b’* stimuli had negative polarity and no cue was delivered (**Fig. 4B**, *Pair*). Following >1.5 hours of stimulation, sensory stimulation within the receptive field was transiently turned off in order to probe neural responses to motor corollary discharge signals in the two contexts; these responses reflect the reafference predictions generated by the ELL’s internal model (**Fig. 4B**, *Probe*).

**Fig. 4.**
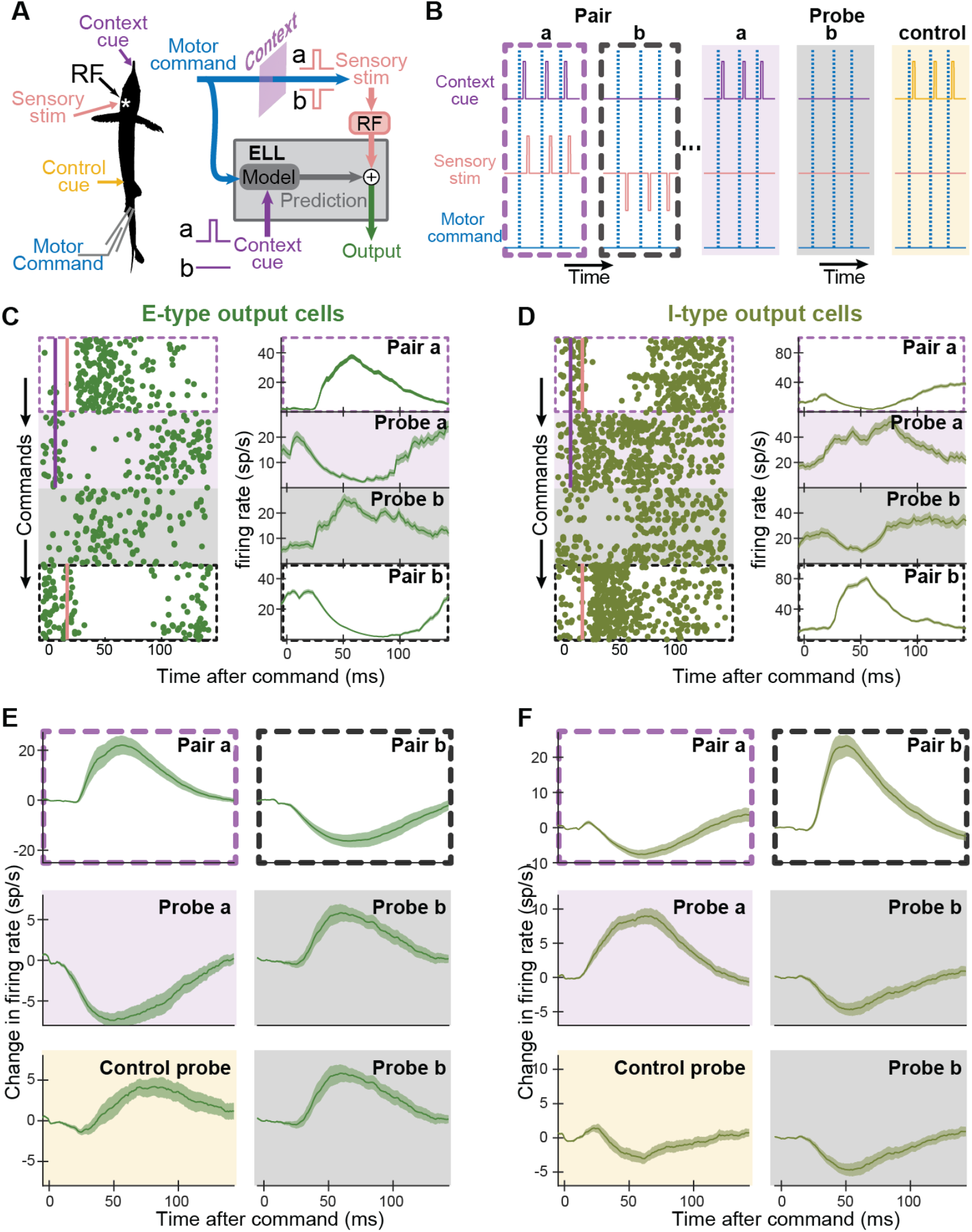
ELL output cells generate context-specific predictions. (**A-B**) Experimental design. EOD motor command recording (blue), local electrosensory stimulus delivered within the RF of recorded neurons (sensory stim, pink) and in two locations outside the RF (context cue, purple; control cue, yellow). Pairing: RF stimuli delivered 11 ms after each EOD command, alternating between two contexts; context a: positive polarity stimulus preceded by a context cue (0.5 ms after command); context b: negative polarity stimulus, no context cue. Probing: transiently omitting RF stimuli and measuring response to the EOD command with and without context cue, and with unpaired control cue to reveal negative images of predicted sensory input. (**C-D**) Command-aligned raster plots (left) and PSTH (right) of example E-type (C) and I-type (D) output units, following >1.5 hours of pairing. Consecutive blocks, from top: Pair a, E-type is excited and I-type is inhibited; Probe a (RF stim removed), E-type inhibited and I-type excited (negative image of Pair a); Probe b (context cue removed), cells switch to opposite response (negative image of Pair b); Pair b (RF stim re-introduced), E-type inhibited and I-type excited. (**E-F**) Summary of results for all recorded E-type (E) (n=21, Pair a-b, Probe a-b; n=13, Control probe) and I-type (F) (n=27, Pair a-b, Probe a-b; n=16, Control probe) output cells.

Recordings were made from ELL output cells (corresponding to the type 2 units described above); these cells can be categorized according to their sensory response into E-type and I-type (similar to ON-and OFF-cells in the visual system). We focused on the zone of the ELL mediating passive electrolocation, where mechanisms for sensory cancellation are most thoroughly understood (*15, 17, 18*). As expected, output cells exhibited opposite polarity spike rate modulations in response to the opposite polarity sensory inputs delivered in the two contexts (E-type output cells were excited in *context a* and inhibited in *context b*, while I-cells exhibited the converse pattern) (**Fig. 4C-F**, *Pair a, Pair b*). Thus, the same EOD motor command was associated with two opposite reafference patterns. This is similar to the situation observed in freely swimming fish, in which the same tail movements evoke opposite changes in reafference depending on the location of the fish in the tank (**Fig.1 F,G**). When the sensory stimulus was turned off in the presence of the context cue (**Fig. 4C-F**, *Probe a*) output cell responses resembled a negative image of the response during pairing in *context* a (**Fig. 4C-F**, *Pair a*). Strikingly, when the context cue was removed, neural responses immediately switched polarity (**Fig. 4C-F**, *Probe b*), i.e. a negative image of the response during pairing in *context b* was observed (**Fig. 4C-F**, *Pair b*). Such context-specific negative images were not observed using a control cue which was located outside of the receptive field but (unlike the context cue) was not previously paired with sensory input delivered to the receptive field (**Fig. 4E,F**, *Control probe*). Additional experiments showed that stimulation of widely-distributed body regions was sufficient for forming context-specific negative images and that stimulation outside of the receptive field had little or no effect on neuronal responses without prior pairing (**Fig. S6**). Similar context-specific negative images were also observed in experiments in which a context cue was used to determine the timing of the local sensory stimulus, rather than its polarity (**Fig. S6**). These results demonstrate that ELL circuitry is capable of combining sensory and motor signals to learn context-specific predictions of self-generated sensory input.

Prior electrophysiological studies in a variety of systems have identified key roles for internal, motor-related signals in predicting and cancelling self-generated sensory input (*30-37*). However, the neural mechanisms for reafference cancellation under naturalistic conditions in freely behaving animals remain poorly understood. Using novel methods for underwater neural recordings in freely swimming electric fish, we provide evidence that reafference cancellation cannot be based purely on motor signals, but additionally requires an estimate of context derived from external sensory cues. These results suggest that the relatively simple, cerebellum-like circuitry of the ELL contains an unexpectedly sophisticated, context-dependent internal model of the environment.

Given its experimental tractability, the ELL may provide opportunities for addressing the challenging question of how internal models are implemented at the synaptic, cellular, and circuit levels. Key questions for future studies are: (1) how a reliable representation of context is extracted from electrosensory input and (2) how context information (which is likely conveyed by electrosensory feedback pathways to granule cells of the ELL and an associated region of the cerebellum (*27, 38, 39*)) can be used for real-time cancellation of sensory reafference despite inevitable processing delays. While the use of so-called forward models to compensate for such processing delays is a proposed function of the cerebellum (*40*), the neural basis for such a function has yet to be elucidated.

## Materials and Methods

### Experimental Model and Subject Details

Adult male and female Mormyrid fish (15-22 cm in length) of the species *Gnathonemus petersii* were used in these experiments. Fish were housed in 60-gallon tanks in groups of 5-20. Water conductivity was maintained between 60-100 microsiemens both in the fish’s home tanks and during experiments. All experiments performed in this study adhere to the American Physiological Society’s *Guiding Principles in the Care and Use of Animals* and were approved by the Institutional Animal Care and Use Committee of Columbia University.

### Experimental Procedures

#### Implant design

The implant included a miniaturized headstage, an electrode interface board (EIB), a microdrive, and a microelectrode probe. The headstage was based on the Open-Ephys design (Open Ephys, https://open-ephys.org) and was modified to have a horizontal alignment during recordings so as to minimize overall implant height. The headstage was encapsulated in epoxy resin to protect it from water damage. The microdrive was based on previously published design (*41*) and was modified to minimize implant size, conductivity, and weight. The EIB was attached to the microdrive screw and stereotrodes were prepared by twisting pairs of 0.0005” tungsten wires (California Fine Wire, Grover Beach, CA) using a dedicated assembly station (Neuralynx, Bozeman, MT). Between 2-8 Individual stereotrodes were then inserted into the microdrive and connected to the EIB using gold pins (Neuralynx, Bozeman, MT). The reference channel was connected to ground, and a 10 cm insulated silver or stainless-steel wire was soldered to the ground channel. Epoxy resin was applied on both sides to the assembled EIB to protect and insulate the wire connections. Vaseline gel was applied to the EIB connector before mating it with its headstage counterpart and applying Cyanoacrylate glue to waterproof the connection. A similar procedure was used to secure and waterproof the headstage-to-tether connector. Lastly, a small Styrofoam piece was attached to the headstage to neutralize the overall buoyancy of the implant. The microdrive was fully retracted and the electrode wires were cut to initial length of 3 mm on the day of implantation.

#### Surgery and Implantation

Fish were anesthetized (MS:222, 1:25,000) and held against a foam pad. Aerated water with MS:222 was continually passed over the fish’s gills for respiration throughout the surgery. Skin on the dorsal surface of the head was removed and a long-lasting local anesthetic (0.75% Bupivacaine) was applied to the wound margins. A hole was drilled in the skull close to the anterior edge of the wound and a single bone-screw, pre-soldered to a 2 cm long silver wire, was installed. A 2 mm diameter hole was then drilled in the posterior portion of the skull overlying the medial zone of ELL (MZ, 1-2 mm lateral to midline). A glass microelectrode filled with 2M NaCl was inserted into the granular layer of ELL (∼3.5 mm depth). To identify the receptive field (RF) of the intended implantation site, short pulses (200 us, 20 uA) were delivered via a silver chloride dipole electrode that was moved across the fish’s skin while observing the evoked local field potential (LFP). Once the RF was identified, the glass microelectrode was removed and the implant was positioned above the craniotomy. The bone screw wire was soldered to the implant ground wire, and then the implant was lowered until the bottom of the microdrive touched the skull surface (inserting electrode ∼2 mm deep). The implant was then cemented in place using Metabond (Parkell, Edgewood, NY), making sure to completely cover the uninsulated bone-screw and wire. Anesthesia was removed, the fish was respirated with aerated water until resumption of a steady EOD rate and then transferred to the experimental tank with a tether connected to a motorized commutator. Fish were allowed to recover for at least 2 hours before electrodes were further advanced.

#### Chronic experiments

The recording chamber was kept at ambient air temperature of 29°C and with a 12 hour daylight cycle. Fish were placed in an 18” diameter circular arena inside a rectangular tank (18.5”X20.5”) that was filled with constantly aerated water to 4” depth; the relatively shallow water eliminated most motion along the vertical axis. The tank was backlit with an infrared LED array. Water temperature, conductivity, pH, and ammonia concentration were monitored daily. Fish were fed with live blackworms (Eastern Aquatics, Lancaster, PA) once a day. Recordings were conducted continually, either in an empty arena or with a vertical brass or plastic pole (0.25” diameter) placed at the center. Electrodes were advanced by lightly sedating the fish with metomidate hydrochloride (Western Chemical Inc, Femdale, WA) in a small chamber placed within the arena and turning the microdrive thumbnut (minimum 125 um between recording sites).

Experiment control and data logging was performed using a custom-made Bonsai workflow (https://bonsai-rx.org/). The tank was video recorded using an IR sensitive camera (FLIR systems, Wilsonville, OR) at 50 fps. Real-time tracking was used to monitor the fish’s heading azimuth, and the commutator was automatically rotated to relieve tether-torsion whenever the fish completed a full turn in either direction. The fish’s EOD rate was monitored in real-time using a pair of carbon electrodes attached to the sides of the arena. The physiological data from the implanted electrodes (30K samples/s) and the 3-axis accelerometer readings (7.5K samples/s) were recorded using an Open-Ephys data-acquisition board (https://open-ephys.org).

#### Acute experiments

Surgery was conducted as detailed above to expose ELL for recording. Gallamine triethiodide (Flaxedil) was given at the end of the surgery (∼20 μg/cm of body length) and the anesthetic was removed. Aerated water was passed over the fish’s gills for respiration. Paralysis blocks the effect of electromotoneurons on the electric organ, preventing the EOD, but the motor command signal that would normally elicit an EOD continues to be emitted by the electromotoneurons at a variable rate of 2 to 5 Hz. This EOD motor command signal was recorded with an Ag-AgCl electrode placed over the electric organ. The command signal lasts about 3 ms and consists of a small negative wave followed by three larger biphasic waves. Onset of EOD command was recorded as the negative peak of the first large biphasic wave in the command signal. Extracellular single-unit recordings were made using glass microelectrodes (2-10 MΩ) filled with 2M NaCl, as described previously (*18, 42, 43*). Recording locations within the ventrolateral zone of the ELL were established using characteristic field potentials evoked by the EOD command along with mutli-unit responses to low-frequency electrosensory stimuli. Recordings were restricted to units with receptive fields on the left side of the face. Ampullary electroreceptor afferents, E cells and I cells are located in different layers of ELL and have distinctive electrophysiological characteristics (*18, 42, 43*). Importantly, E cells are excited by the same stimulus polarity as afferents while I cells are excited by the opposite polarity. Previous studies using intracellular recording and biocytin labeling and antidromic stimulation from the midbrain have shown that E and I cells correspond to two morphologically distinct types of ELL efferent cells known as large fusiform and large ganglion cells (*28*).

Local sensory stimuli (monophasic square pulses 200 us width, ±1 uA) were delivered between a patch of carbon placed adjacent to the left side of the fish’s face and an Ag-AgCl wire placed in the stomach of the fish at an 11 ms delay after the EOD motor command. The results presented in **Fig. 4** were obtained using the following ‘polarity context’ protocol. In ‘context a’, the local stimulus had positive polarity (i.e., excited E cells and inhibited I cells) and a second local stimulus (the Context cue) was delivered via an Ag-AgCl dipole located either next to the fish’s chin (schnauzenorgan) or to the fish’s trunk (monophasic square pulses 200 us width, 25 uA, delivered 0.5 ms following EOD onset); in ‘context b’, local mimic polarity was negative (i.e., inhibited E cells and excited I cells) and no preceding context cue was given. Negative images were probed by transiently turning off the local EOD mimic at the face to observe the neural response in the absence of afferent input (*15*). In another set of experiments, we used a ‘temporal context’ protocol, in which polarity was positive in both contexts, but in context a mimics were delivered 65 ms after the EOD command signal, while in context b they were delivered 4.5 ms after the command (see **Fig. S6**).

### Quantification and Statistical Analysis

Unless stated otherwise, all analyses were performed using custom Matlab code (Mathworks, Natick, MA).

#### Pose Tracking

Video frames were down-sampled to 400×416 pixel resolution and then analyzed using DeepLabCut (*44*) to extract the coordinates of 9 features (tip of chin, mouth, two points on trunk, two points on tail, caudal fin, left and right tips of pectoral fins). The locations of these features in every frame were converted into head center coordinates, heading azimuth, and 7 pose angles. The fish’s tail position was extracted using the first Principal Component of the body angles (excluding the pectoral fin angles).

#### Spike sorting

EOD artifacts were detected using threshold crossing of the first derivative of the electrode voltage; a time window 0.5 ms prior to 1 ms post EOD artifacts was then removed. Low-frequency LFP signals were removed by subtracting a median-filtered (2 ms) version of the raw signal. Spike detection and sorting was performed using KiloSort (*45*) and then manually curated using Phy2 (https://phy.readthedocs.io/).

#### LFP processing

To obtain the LFP amplitudes associated with each EOD, voltage traces 1-15 ms after every EOD were extracted for all channels. These traces were high-pass filtered (100 Hz cutoff) and the peak-to-peak amplitude was measured for each channel. Finally, the LFP amplitude data from each channel were z-scored, and the values obtained from the two channels of each stereotrode were averaged.

#### EOD Data Collection

For each EOD recorded, the following data were collected: instantaneous discharge rate (i.e., reciprocal of preceding inter-EOD time interval); z-scored LFP amplitudes of all stereotrodes (see above); spike count in the time window 1-25 ms following the EOD for each sorted single unit; head-center coordinates, heading azimuth, and 7 body angles (extracted from video tracking, see above), interpolated to the EOD onset time; 3-axis accelerometer readings, interpolated to the EOD onset time.

#### Exafference and reafference maps

In order to compute the exafference and reafference maps of a physiological variable (LFP amplitude or spike count), we first transformed the allocentric fish position (head coordinated + heading azimuth) into egocentric (head centered) wall or object position; wall position was defined as the distance and direction of the closest point on the wall to the head center (i.e. a line between this point on the wall and the tank center passes through the fish’s head coordinates). The converted wall/object coordinates (*x, y*) were binned using a two-dimensional grid (N=25 bins in each dimension). In each bin, linear regression was performed to fit the relationship between the physiological variable (*r*, the ‘response’) and a zero-centered ‘input’ variable *t* (tail position or LFP amplitude):

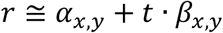

The ordinate intercept of the fit (*α*_*x,y*_) is the average response given an average input, and hence the input-controlled response modulation across all object locations (i.e., the exafference). The slope *β*_*x,y*_ provides the response modulations due to *t* (i.e., the reafference due to *t*). The overall tail/LFP modulation across the entire map was defined as the average absolute value of the slope:

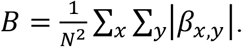

The distribution of *α, p*_*α*’_, was computed using a N=25 bin histogram and the exafference Spatial Selectivity was defined as:

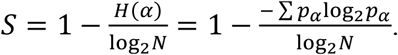

Hence, *S* = 0 for a completely uniform distribution (i.e., a graded map with equal representation to all response levels), and *S* ≈ 1 for a map with a single over-represented ‘background’ value and just a few bins diverging from that value (the ‘receptive field’ of the response).

#### Electrostatic Multipole Model

To model the fish’s electric field and its modulations due to tail motion and external objects, we used the multipole electrostatic model developed by Chen et al (*19*). To simplify the analysis, we assumed cylindrical symmetry that eliminated modulations on the vertical axis. The fish was modeled as a collection of 340 point charges distributed along the main body axis (20 poles/cm along 17 cm total fish length), each charge at position 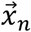.The charge of the most caudal pole was *q*_0_ = −1, while that of all other N=339 poles was *q*_*n*_ = 1/*N, n* ∈ [1: *N*], so that the overall charge was 0. The electric potential *ϕ* generated by these poles at some position 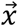 was therefore:

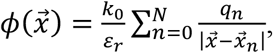

where *k*_0_ is Coulomb’s constant and *ε*_*r*_ the relative permittivity of water. The wall effect of a circular arena with radius *R*=23 cm was approximated using the image method (*46*), where for each charge *q* positioned at distance *d* from the arena’s center, an image with charge *q*^′^ = *α*_*k*_*q* at position *d*^′^ = *R*^2^/*d* was added; 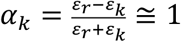 where *ε*_*k*_ is the permittivity of the wall. The electric field 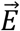 due to all poles (real and imaged) at position 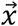, was therefore:

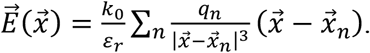

The transdermal potential *z* at a receptive field located in some position 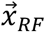 on the fish’s skin was assumed to be proportional to the normal component of the electric field at that position:

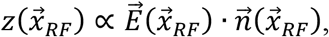

where 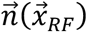 is the normal unity vector (i.e. the direction perpendicular to the skin) at the receptive field. The fish’s skin shape was modeled as a semicircular head (1.5 cm diameter) and an isosceles triangular body extending to the tip of the tail. The rostral 45% of the body (7.65 cm) was maintained straight, while the caudal 65% of the body (11.05 cm) was rotated to imitate tail bending (bending range ± *π*/3 rad).

#### ELL Computational Model

The data for the model was generated using the electrostatic multipole model (see above); transdermal potential was calculated on a grid of wall distances and directions (*r, φ*, 35×36 grid), for 25 different tail angles (*t*_0_, in the range -0.3π to 0.3π), and at 25 different RFs on the model fish’s head:

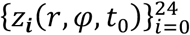

The neural network model approximated the known cerebellum-like architecture of the ELL, and simulated reafference cancelation using an internal model. The input to the internal model (‘mossy fiber’ inputs), 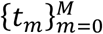, included combinations of the motor corollary discharge (tail position, *t*_0_), the wall distance and direction (explicit context, *r, φ*), the transdermal potential from all 25 RFs 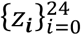, and the unity (bias) constant. This input drove a layer of K=1000 Granule Cells (GC) with a hyperbolic tangent transfer function, via a random, non-plastic, sparse (1-3 connections per GC) weight matrix ***β***^*GC*^. The outputs of the GCs converged onto a single output neuron with a weight vector ***w*** trained using error gradient descent. Note that the expansive GC layer of this model enables high performance without requiring a biologically implausible backpropagation algorithm (*47*). The analyses were also repeated using a standard backpropagation neural network (100 neurons in the hidden layer) producing similar results.

The network was trained to predict reafference using supervised learning. To produce the training signal, we used the sensory signal obtained with a straight tail as the exafference component:

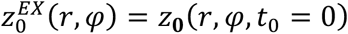

The target signal for training was the difference between the exafference component and the simulated sensory input:

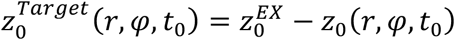

The model is trained to predict this target signal from the GC inputs:

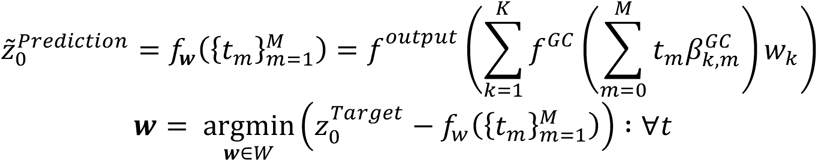

where *f*_*w*_ is the model’s total transfer function and ***w*** is the output layer’s weight matrix. Finally, the output (filtered) sensory signal is given by:

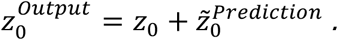

## Acknowledgments

We thank L.F. Abbott, D.M. Wolpert, E. Ahissar, S.Z. Muller, and F. Pedraja for discussions and comments on the manuscript.

## Funding

National Institutes of Health grant NS075023 (NBS)

National Institutes of Health grant NS118448 (NBS)

Irma T. Hirschl Trust (NBS)

## Author contributions

Conceptualization: AW, NBS

Methodology: AW, NBS

Formal analysis: AW

Investigation: AW, NBS

Visualization: AW

Funding acquisition: NBS

Project administration: NBS

Supervision: NBS

Writing – original draft: AW, NBS

Writing – review & editing: AW, NBS

## Data and materials availability

Experimental data files are available at the Columbia University Academic Commons repository.

## Supplementary Materials

Figs. S1 to S6

Movies S1 to S3

