## Supplemental Material for "An Internal Model of Sensorimotor Context in Freely Swimming Electric Fish"

**This file includes:**

Figs. S1 to S6
Captions for Movies S1 to S3

**Other Supplementary Materials for this manuscript include the following:**

Movies S1 to S3

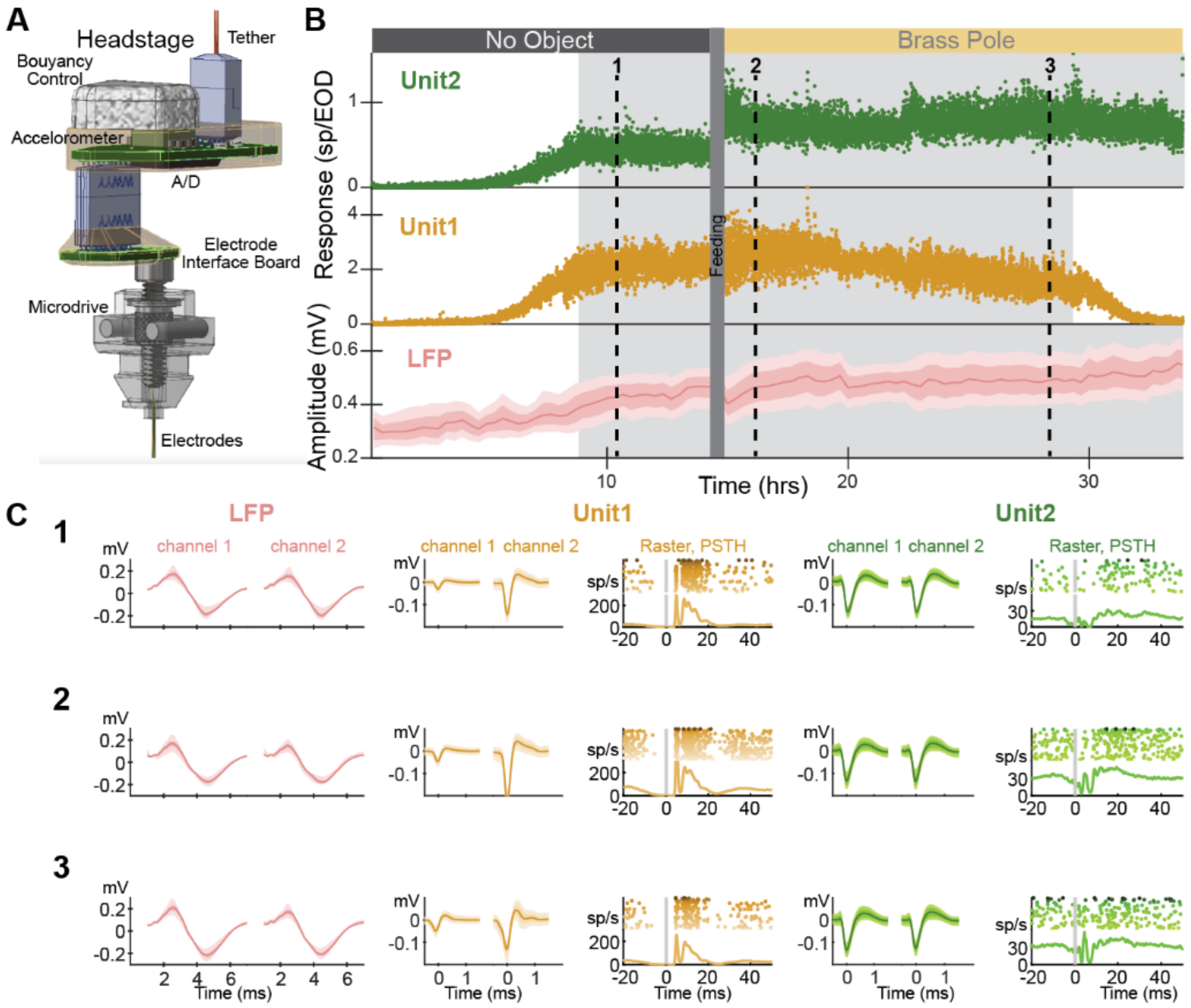


**Fig. S1. Long-term electrophysiological recordings in freely swimming electric fish** (**A**) Implant design, including 3D-printed microdrive, electrode interface board and headstage. All electronics are embedded in water-proof epoxy. Styrofoam block is used to neutralize buoyancy. (**B**) Example of continuous recording from one site in ELL for over 33 hours, which included two recording segments: 13 hours with no object and 20 hours with brass pole positioned at center of tank. Top: response of two example units (spikes/EOD, smoothed and down-sampled by a factor of 100); bottom: amplitude distribution of LFP from one stereotrode (mean, bold line; dark shading, IQR; light shading, 10-90% range). Analyses were performed in each recording segment separately, using only epochs in which activity was stable (grey shading). Note the three timepoints marked 1-3. (**C**) Left: LFP waveforms at the three timepoints marked in B. Middle: Spike waveforms (left) and EOD responses (EOD-aligned raster and PSTH) for unit1 at the three timepoints in B. Right: Spike waveforms (left) and EOD responses (EOD-aligned raster and PSTH) for unit2 at the three timepoints in (B) (in all waveforms: mean, bold line; dark shading, IQR; light shading, 10-90% range).

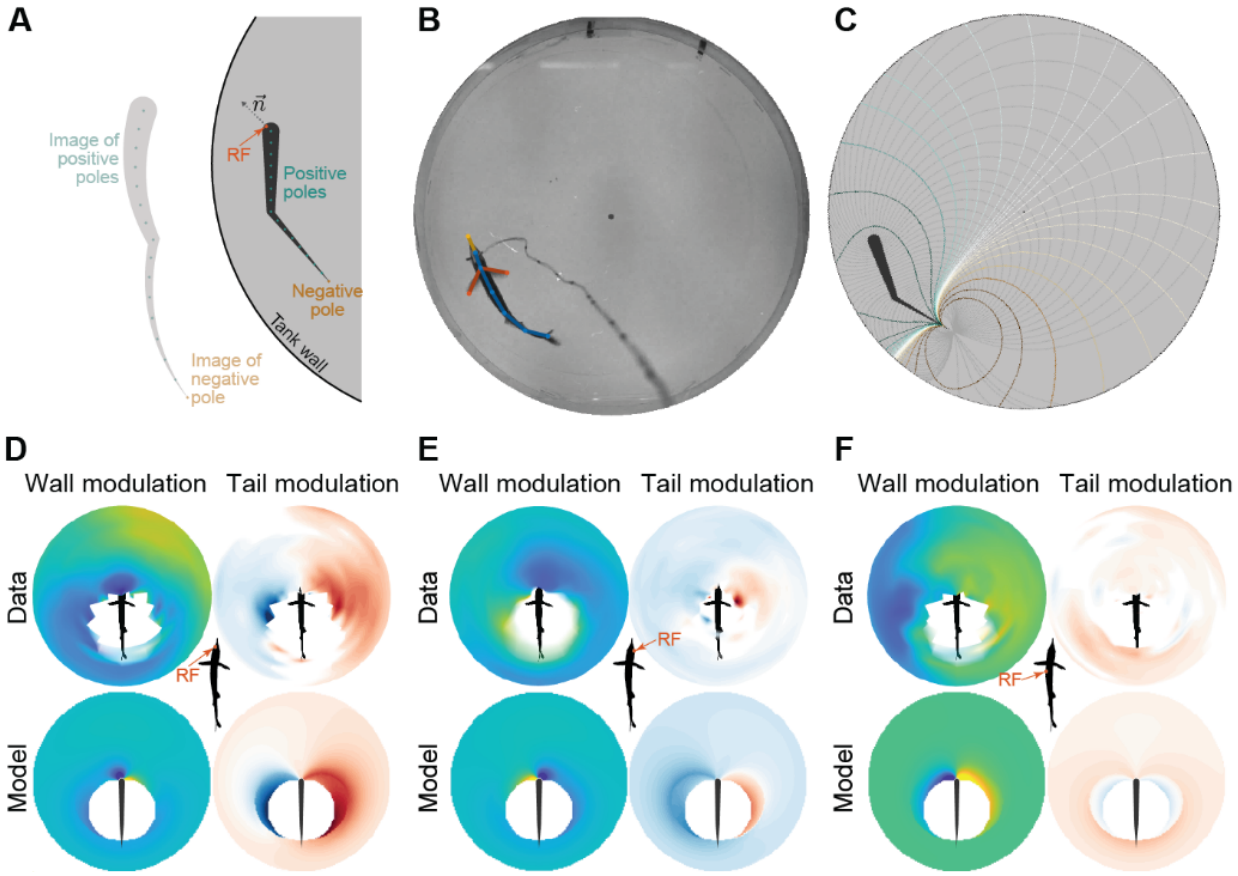


**Fig. S2. Electrostatic multipole model recreates LFP exafference and reafference across the somatotopic map** The model used here is simplistic in many respects. First, it assumes *cylindrical symmetry*, i.e. it ignores the boundary effects of the water surface and the tank floor. These effects were approximately constant, as the shallow water-level in these experiments (10 cm depth) minimized vertical motion. However, the model is ill-suited to ventral or dorsal receptive fields (RFs), as well as to investigate the effects of roll and pitch movement. A second simplification of the model is that it ignores the boundary effects of the fish's skin. Nevertheless, this simple model successfully captures qualitative exafference and reafference patterns obtained from a variety of RFs (see D-F). (**A**) The electric fish is modeled as a collection of small positive point charges along its main body axis (20 poles/cm, only 1-in-20 is presented, green) and a single large negative point charge at the caudal end of the electric organ (brown); overall charge is neutral. The non-conductive tank wall is equivalent to image poles distributed outside (note the distortion due to the wall curvature). The resulting electric potential and field can be computed at any point in the tank using Coulomb's law. The transdermal potential at some RF on the skin is proportional to the electric field's component normal to the skin (marked $\vec{n}$) at that RF. (**B**) Snapshot of implanted fish from overhead camera, with tracked features and skeleton model used to derive fish posture (yellow, chin; blue, six points along main body axis; red, pectoral fins). (**C**) Derivation of electric potential (colored lines) and field (grey lines) for implanted fish using the electrostatic model. (**D-F**) Wall modulation (exafference, regression intercept, left) and tail modulation (reafference, regression slope, right) maps derived from LFP amplitude data (top row) and electrostatic multipole model (bottom row) for various RFs ((D) left base of chin, (E) right front face, (F) left mid-trunk).

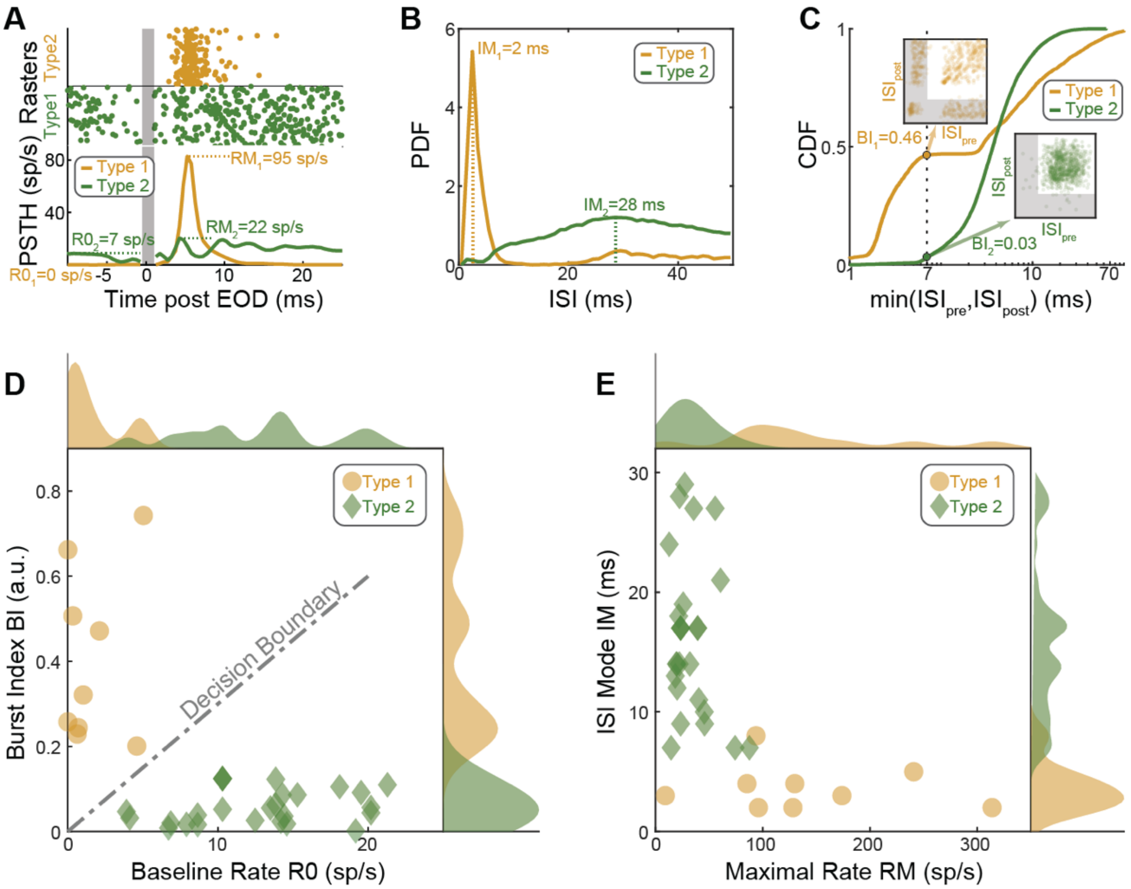


**Fig. S3. ELL single-unit categorization.** (**A**) EOD-triggered raster and peri-stimulus time histogram (PSTH) plots for two example ELL units. For each unit, the baseline firing R0 (firing rate in the time window 5-20 ms prior to EOD, averaged across all EODs preceded by an interval of at least 50 ms) and the maximal firing rate RM (maximal value of PSTH, computed using 1 ms time-bins) were calculated. (**B**) Inter-spike interval (ISI) probability distributions (PDF) for the two example ELL units presented in A. For each unit, the ISI mode IM (most frequent value after rounding to 1 ms resolution) was calculated. **(C**) Cumulative distributions (CDF) of the smaller of the ISIs preceding and following each spike, for the two example neurons in A and B (log-scale). The burst index BI was defined as the proportion of spikes either preceded or followed by an ISI smaller than 7 ms. Insets: ISI first-return maps (plot of the ISI following vs. the ISI preceding each spike). Note the 4-cluster distribution typical of bursty neurons for Type 1 example. Shaded areas in insets depict spikes under the burst threshold (7 ms). (**D**) Distribution of Baseline Rate R0 and Burst Index BI for all ELL units. Note the decision line used to discriminate between Type 1 (yellow circles, low R0 and high BI) and Type 2 (green diamonds, high R0 and low BI). PDFs of each variable are shown at the margins. (**E**) Distribution of Maximal Rate RM and ISI Mode IM for all ELL units. Type 1 (yellow circles) had high RM and low IM, while Type 2 (green diamonds) had low RM and high IM. PDFs of each variable are shown at the margins.

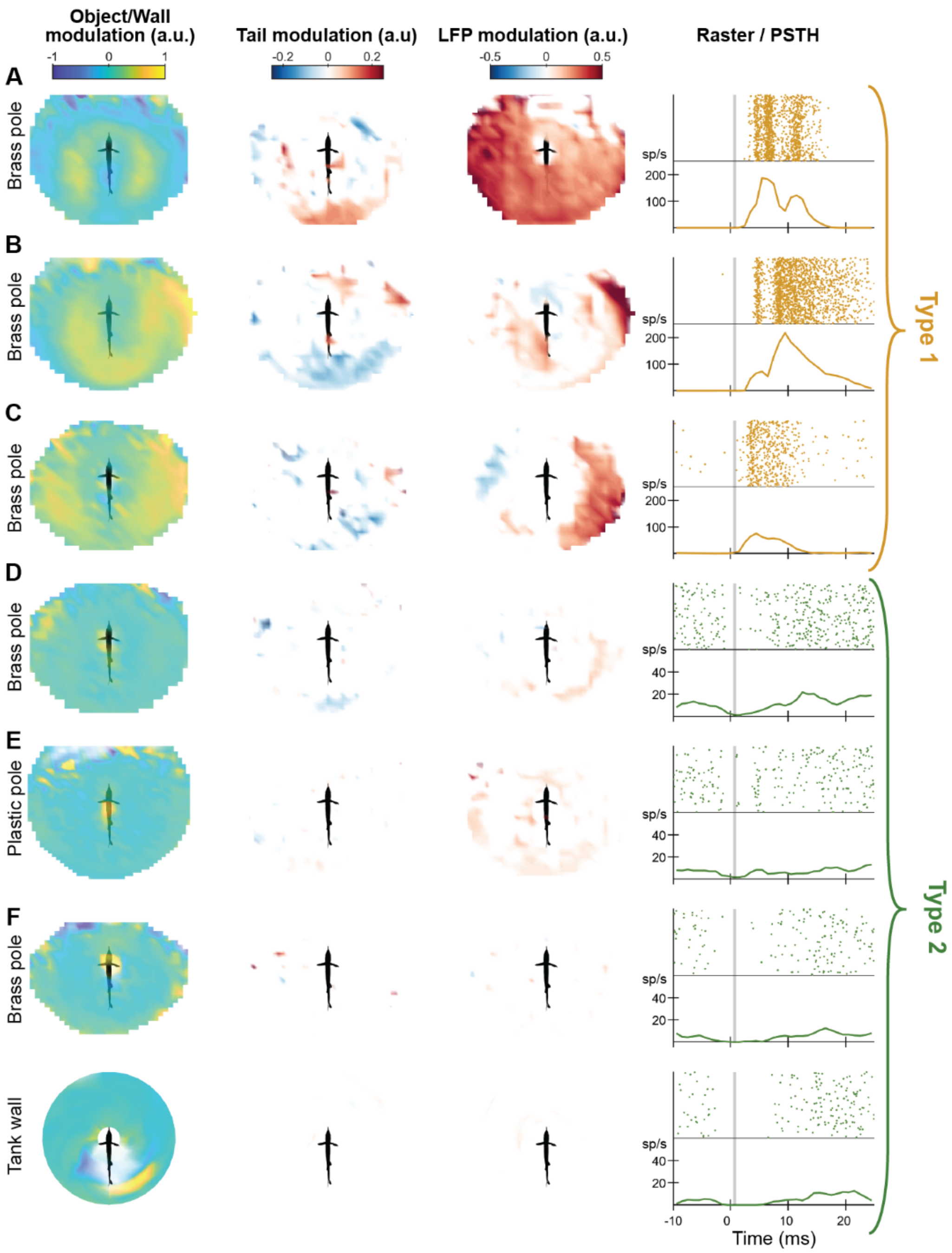


**Fig. S4. Examples of ELL single-units.** From left: Object/wall modulation map (tail-regression intercept, z-score) for example units; Tail modulation (tail-regression slope, z-score/z-score); LFP modulation (LFP-regression slope, z-score/z-score); Raster plot (top) and PSTH (bottom, note scale-difference between Type 1 and Type 2 units). (**A-C**) Type 1 units exhibited spatially diffuse responses to the brass pole and substantial reafference. (**D-F**) Type 2 units exhibited spatially selective responses to objects and negligible reafference. (D) RF center on the anterior right side of face (brass pole); (E) RF center on the left trunk, response to plastic pole (suggesting an I-type output cell); (F) RF center on the posterior left side of face, top row – unit's response to brass pole, bottom row – same unit's response to wall.

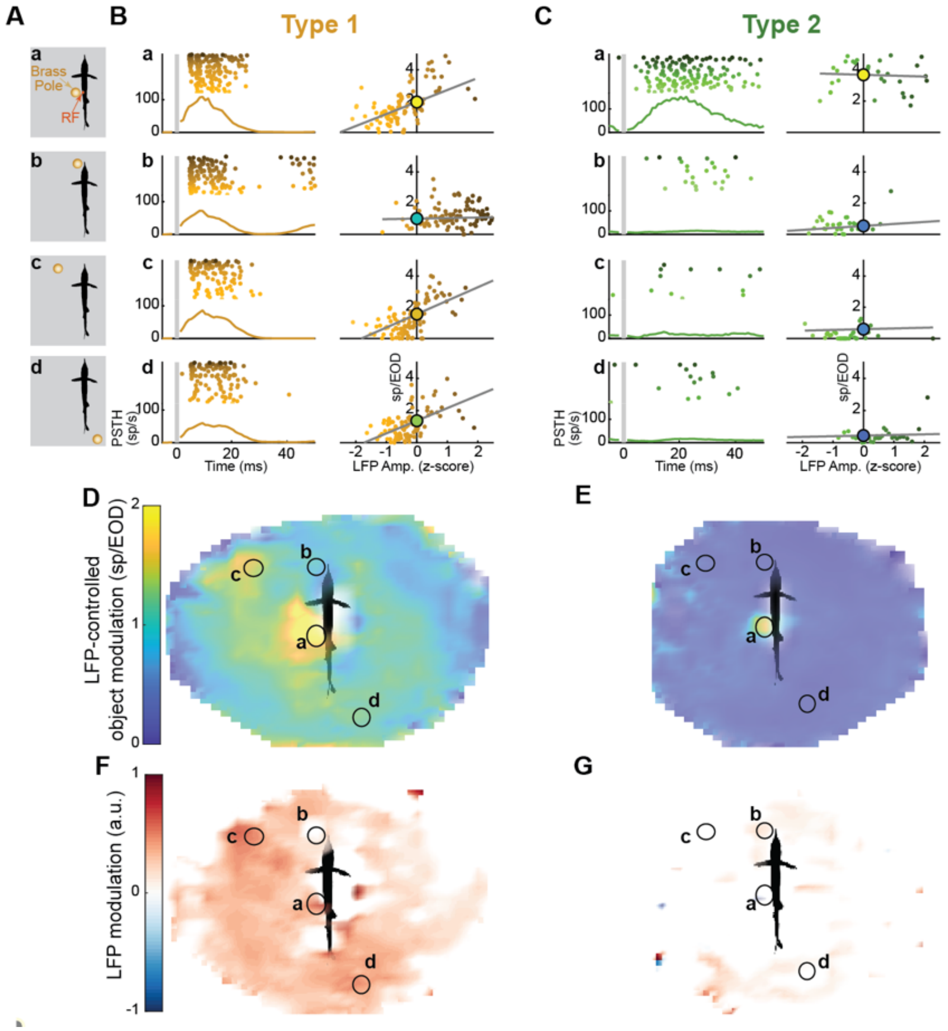


**Fig. S5. Analysis of ELL input-output transformations.** Analysis of the transformation of sensory input into ELL neuronal responses, performed by correlating ELL unit spiking and the LFP amplitude recorded on the same electrode, for all relative positions of an object (vertical brass pole located at the center of the tank). For any given object position, exafference is approximately fixed, and therefore variations in sensory input (LFP amplitude) reflect reafference (from all motor variables). (**A**) Four examples of object position relative to fish. Receptive fields (RF) of the units shown here is on the left trunk. (**B**) Left: EOD-aligned peri-stimuli time histograms (PSTH) and rasters (each ordered by ascending LFP amplitude from bottom to top) of example type 1 unit, for the different object positions. Right: spike count (within 40 ms after EOD, smoothed using boxcar averaging 5 EODs wide) vs. LFP amplitude (z-scored); grey line: linear regression, round marker: regression intercept. For type 1 units, intercept is graded and almost in all object positions reafference (slope) is substantial (positions a, c and d). (**C**) Same analysis as in B for example type 2 (putative output) cell. For these units, intercept is very spatially selective (strong response only for position a, i.e. when object is adjacent to RF), and reafference is cancelled (negligible slope). Importantly, since the bias reflects the unit's response for an average LFP amplitude (i.e. when variations in sensory input across positions are controlled for), the observed spatial selectivity of type 2 units does not arise from a simple non-linear transformation of the sensory input. (**D-E**) LFP intercept (LFP-controlled object modulation) maps for the two units. Note the 4 example positions (labeled black circles). (**F-G**) LFP modulation map (regression slope) for the two units.

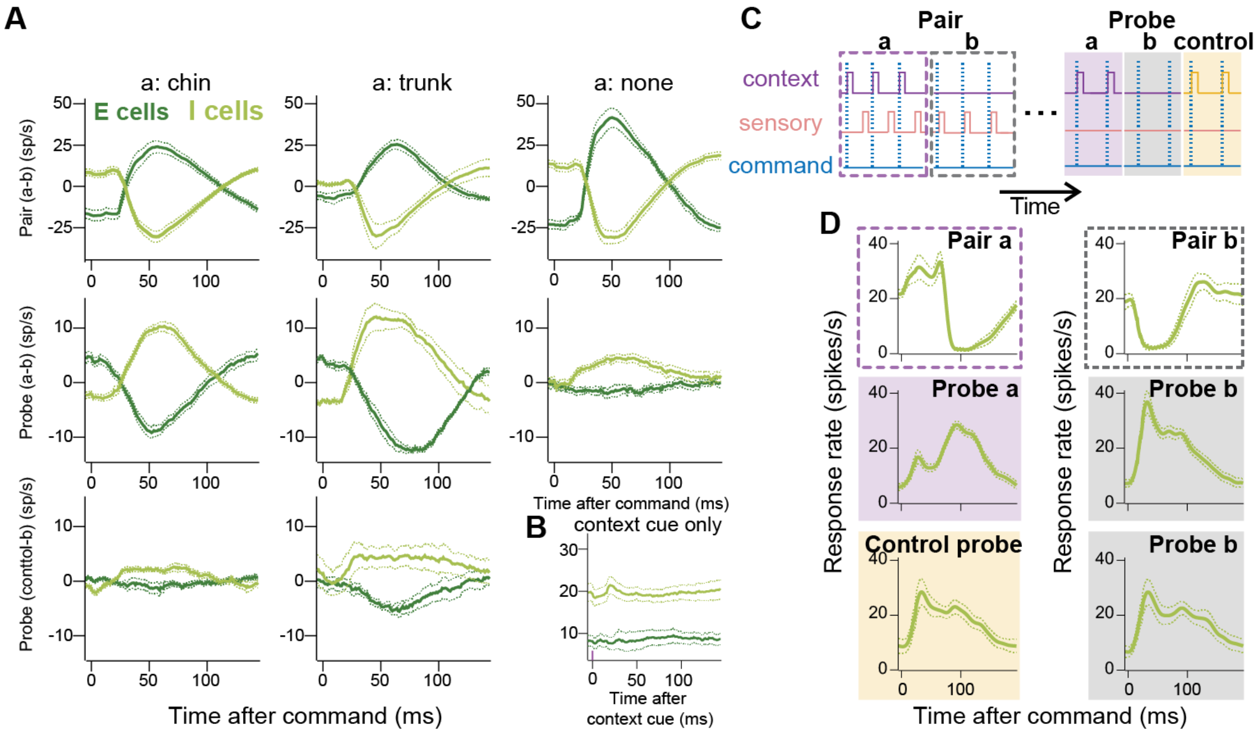


**Fig. S6. ELL output cells generate context-specific predictions.** (**A**) Additional analysis of data presented in **Fig. 4**. Traces show the firing rate difference in the two contexts for all recorded E- and I-type cells. Left column: paired context cue was delivered to the chin (Schnauzenorgan); center column: paired context cue was delivered to the trunk; right column: no context cue was delivered during pairing (control). Top row: response difference between contexts during pairing (a:chin n= 14, E cells, n=19, I cells; a:trunk n=7, E cells, n=8, I cells; a:none n=6, E cells, n=14, I cells); middle row: response difference between context during probing (a:chin n=14, E cells, n=19, I cells; a:trunk n= 7, E cells, n=8, I cells; a:none n=6, E cells, n=14, I cells); bottom row: response difference using unpaired control stim (a:chin n=8, E cells, n=10, I cells; a:trunk n=5, E cells, n=6, I cells). (**B**) Response to the context stimulus alone in the absence of prior pairing with sensory stimulation delivered to the RF (n=4, E cells; n=14, I cells). (**C**) Experimental design for a second series of experiments in which the timing relative to the commands (rather than the polarity) of the local sensory stimulus was dependent on context (See **Methods**). (**D**) Top: average response of ELL I-cells (n=8) during pairing in context a (left) and context b (right). Note the difference in the timing of stimulus-induced inhibition. Middle: negative images revealed during probing in context a (left) and context b (right) (n=8). Note the difference in the timing of the peak response in the two contexts. Bottom: responses to a previously unpaired context stimulus (left; Control probe) and no context stimulus (context b, right) are identical (n=7).

**Movie S1. Long-term behavioral and physiological recordings in freely swimming electric fish.**

**Left**: overhead videography of implanted fish in a tank with a brass pole (played at 12.5 frames/s, i.e. 4X slower than real-time). Tracked body features (circles) and skeleton model (line segments) are overlaid (main body axis, blue; pectoral fins, yellow). **Middle**: dynamics of data variables. From bottom to top: head azimuth, head coordinates, left and right pectoral fin angles, pitch and roll angles (from accelerometer), tail position, EOD instantaneous rate, LFP peak-to-peak amplitude, unit 1 and 2 spike count (in 25 ms post EOD window). **Right**: EOD-triggered electrode voltage trace (bottom), LFP activity (color coded, top left), and units' activity (raster plot, top right).

**Movie S2. ELL wall (exafference) and tail (reafference) modulation maps: experiment and model.**

**Top row**: experimental results. **Bottom row**: electrostatic multipole model results. **Leftmost column**: example video showing implanted fish swimming in tank (no center object, 0.5X speed). Red circle: position of closest point on wall to fish's head (a radius line ending in this point passes through the fish's head). Top-left inset: LFP traces from one implanted electrode with RF on the left side of the face (mV/ms). Bottom panel shows same video with overlaid electric potential (colored lines) and field (grey line), derived from multipole model. **Second column**: average LFP amplitude as function of wall relative position (distance and direction), plotted for different tail positions. **Third column**: linear regression intercept map (wall modulation, exafference). **Rightmost column**: linear regression slope map (tail modulation, reafference).

**Movie S3. ELL object (brass pole, exafference) and tail (reafference) modulation maps examples.**

**Top**: Average LFP amplitude (electrosensory input, left) and single units' response (spike count within 25 ms window post EOD, middle and right) across object positions, for different tail positions. **Center**: Object modulation map (regression intercept, exafference). **Bottom**: Tail modulation map (regression slope, reafference). Electrosensory input and type 1 cells (putative interneurons) exhibit a spatially diffuse response pattern that is strongly modulated by tail motion; tail modulation is spatially patterned (context dependent). In contrast, type 2 (putative output) cells exhibit a highly spatially selective response that is invariant to tail motion regardless of external context.
